# Differential Ubiquitination of Host Proteins in Response to IPNV and ISAV viruses in Atlantic salmon

**DOI:** 10.1101/2024.11.05.622028

**Authors:** Robert Stewart, Xoel Souto Guitián, Ophélie Gervais, Yehwa Jin, Sarah Salisbury, Samuel A. M. Martin, Maeve Ballantyne, Beatriz Orosa-Puente, Diego Robledo

**Affiliations:** The University of Edinburgh, Easter Bush Campus, Midlothian EH25 9RG, UK; University of Santiago de Compostela, Santiago de Compostela 15705, Spain; Centre Scientifique de Monaco, 8 Quai Antoine 1er, 98000, Monaco; The Center for Aquaculture Technologies, 8445 Camino Santa Fe, Suite 104, San Diego, CA 92121, USA; Scottish Fish Immunology Research Centre, School of Biological Sciences, University of Aberdeen, AB24 2TZ, UK

**Keywords:** Fish, aquaculture, immune response, ubiquitin, virus, proteomics, RNA sequencing

## Abstract

Viral diseases remain a major barrier to the sustainable production of farmed fish, primarily attributable to the absence of effective prevention and treatment options. Understanding host-pathogen interactions can guide the development of vaccines, antiviral therapies, or gene editing strategies. Ubiquitination is a key cell signalling molecule, known to regulate many aspects of immune functions but currently understudied in fish. This study leverages ubiquitin-enriched mass spectroscopy complemented with RNA sequencing to characterize the role of ubiquitination in response to infection.

A challenge experiment was conducted by infecting Atlantic salmon head kidney (SHK-1) cells with Infectious salmon anaemia virus (ISAV) and Infectious pancreatic necrosis virus (IPNV). At 24- and 48 hours post-infection dramatic changes were observed in the global ubiquitination state of host proteins. Many post-translational modifying proteins increased in abundance upon ISAV infection, whilst IPNV infection resulted in a reduction in abundance of many of these proteins. Transcriptomics showed a delay in the activation of the antiviral response to ISAV infection, with major upregulation of genes associated with immune pathways only at 48h. On the contrary, IPNV infection resulted in upregulation of classic innate immune response genes at both timepoints. Clear activation of Rig-like receptor pathways is demonstrated in both infections, in addition to upregulation of both conserved and novel antiviral TRIM E3 ubiquitin ligate genes. Network analysis identified clusters of immune genes and putatively regulatory proteins showing differential ubiquitination upon viral infection.

This study highlights the importance of post-translational control of the host innate immune response to viruses in Atlantic salmon. Clear differences in ubiquitination between two viruses indicate either virus-specific post-translational regulation or viral antagonism of the immune response. Additionally, the ubiquitination of various proteins was linked to the regulation of innate immune pathways, suggesting a direct role of ubiquitination in the regulation of antiviral responses.

**AUTHOR SUMMARY:** Ubiquitination is a main cellular regulatory mechanism in all animal species, controlling the activity of many proteins within the cell. The role of ubiquitination in the regulation of immunity is well-established, and several direct interactions with viruses have been described, either as part of a strategy of the host to fight the infection or as a viral mechanism to evade the host immune response. In aquaculture, viral diseases currently represent the most important threat to the sustainability of the industry. However, very little is known about the interplay between ubiquitination and viral pathogens in fish species. In our study, we have assessed the ubiquitination response of Atlantic salmon to two production-relevant viruses, Infectious salmon anaemia virus (ISAV) and Infectious pancreatic necrosis virus (IPNV). We found remarkable differences in the ubiquitination profiles between the two infections, suggesting a key role of ubiquitination on the early immune response. We also discovered an association between the ubiquitination of certain regulatory proteins and the activation of antiviral pathways. This information could help develop new strategies to tackle viral diseases in aquaculture, such as the development of more effective vaccines or antiviral therapies, or inform gene editing efforts to generate disease resistant fish.

## INTRODUCTION

### Aquaculture and disease

Aquaculture is a pivotal tool for meeting the increasing demand for high-quality, healthy protein for human consumption. The UN predicts that food demand will double by 2050 as the global population reaches almost 10 billion^1-3^. In 2020, 17% of global animal-source protein consumed was derived from combined aquaculture and fisheries, however, opportunities to sustainably increase wild fish harvest are limited^4,5^. Therefore, it is anticipated that the aquaculture sector will grow by 36-74% over the next 25 years to reflect the ever-increasing demand for protein^5^. While aquaculture food production has multiple advantages over conventional, land-based animal protein production systems including greater food conversion efficiency^6^, low carbon footprint^7^ and lower competitive land use^8^, the growth of the sector is limited by infectious diseases - one of the leading causes of production losses in fish farming^9^.

### Viral diseases in salmon aquaculture

Despite leading to cumulative high levels of mortality in marine-raised Atlantic salmon ^10^, there are few effective prophylactic and therapeutic options for viral diseases affecting salmonids^11-13^. In particular, Infectious Salmon Anaemia Virus (ISAV) and Infectious Pancreatic Necrosis Virus (IPNV) can have catastrophic impacts on the aquaculture sector, yet have limited prevention and treatment options. In aquaculture, Atlantic salmon are commonly vaccinated against ISAV in many production countries, however, current vaccines do not yet offer complete protection^14^. Additionally, there is currently no vaccine available for IPNV^15,16^. Further insights into the host-pathogen interaction of these viruses have the potential to aid in the development of more effective vaccines and help identify targets for selective breeding and gene editing to improve host disease resistance.

### Ubiquitin in the immune response

Post-translational modifications are key in the regulation of both the innate and adaptive immune response^17,18^. In particular, ubiquitination – the addition to a protein of ubiquitin, a small 76 amino acid peptide highly conserved across eukaryotes^19^ - has a broad range of functional effects on the substrate protein, including changes in activity, stability, compartmentalization and conformation. This dynamic and complex modification can lead to the activation of immune pathways; as seen by the ubiquitin-dependant activation of RIG-I upon sensing of dsRNA to trigger IFN signalling^20^, deactivation by targeting regulatory proteins for proteasomal degradation^21^, or direct antiviral mechanisms such as the blocking of viral entry by the Ubiquitin ligase TRIM5^22^. Due to the co-evolution of host and virus, ubiquitin and the ubiquitin proteasome system (UPS) can have both antiviral and pro-viral roles; for example, during Dengue virus infection, ubiquitination is harnessed for viral uncoating^23^, and SARS-CoV-2 has been shown to hijack its hosts UPS to inhibit STAT2 mediated interferon response^24^.

### Summary and aim

Immune regulation by ubiquitination and ubiquitin-like modifications are crucial for a balanced response to pathogens. In model species, the role of ubiquitination has been demonstrated in both the activation and deactivation of both the innate and adaptive immune response^25^. Teleost fish have a greater reliance on the innate immune response than mammals^26^, hence post-translational modifications may play a more pivotal role in host-pathogen interactions in fish. This research has focused on understanding the role of ubiquitination in response to viral infections (ISAV and IPNV) in Atlantic salmon. Our results reveal clear virus-specific changes in ubiquitination patterns in a salmonid cell line, which can be connected to changes in immune-related pathways at both the proteomic and transcriptomic levels.

## RESULTS

### Global ubiquitination changes in response to viral infection

Salmon head kidney cells were infected with ISAV and IPNV, with cell lysate collected at 24 and 48 hours post-infection. To determine the global host ubiquitin response to viral infection, western blot analysis of whole cell lysates was performed on ISAV and IPNV infected cell cultures at 24 and 48 hours post-infection (Figure 1 A & B). Pan-linkage specific ubiquitin blotting revealed a clear increase in ubiquitination of host proteins for ISAV infection, relative to uninfected, time-matched controls. In contrast, IPNV infection reduced the total amount of ubiquitinated proteins.

**Figure 1:**
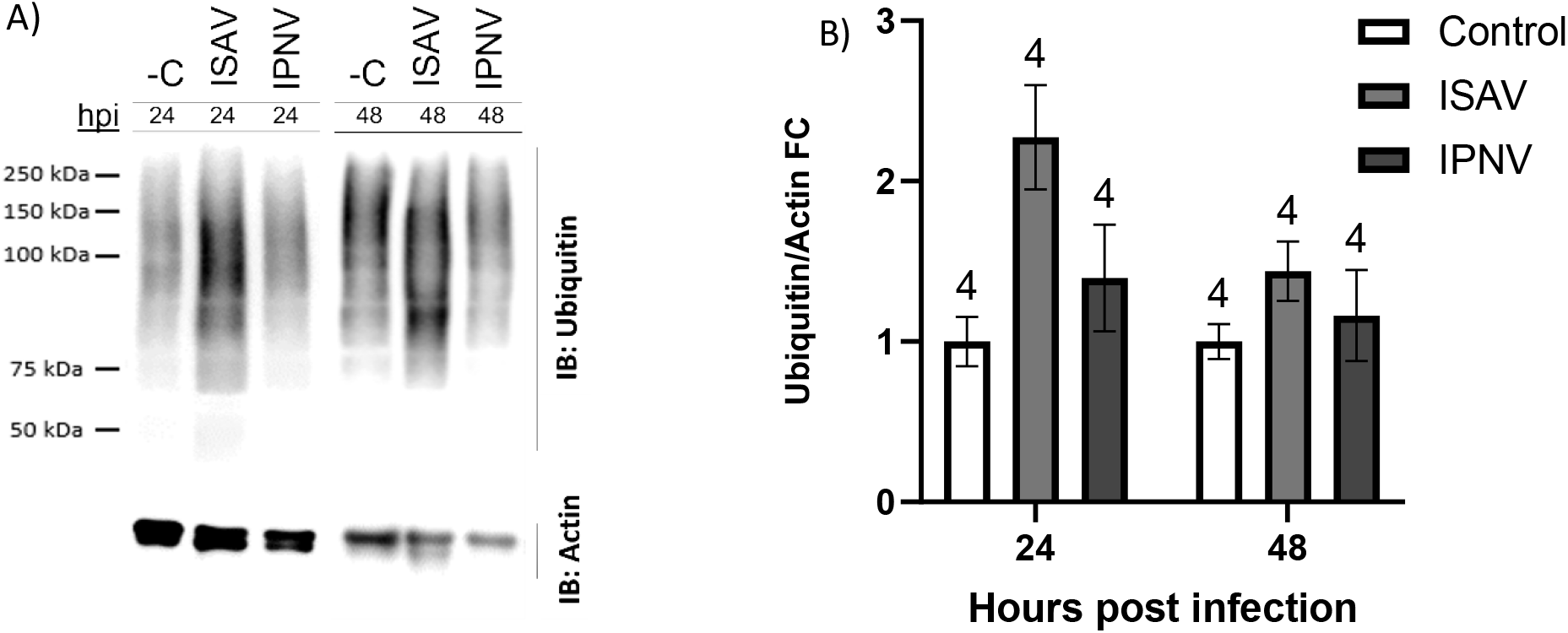
Global ubiquitination state of infected cells. (A) Representative blots of Anti-Ubiquitin western blots from ISAV and IPNV infected whole cell lysate at 24 hpi and 48 hpi. C-denotes negative control (uninfected, time-matched samples). IB: refers to antibodies used for immune-blot. (B) Quantification of western blot bands. Fold change relative to time matched control. Errors bars represent SEM. Sample size is displayed above bars.

### Ubiquitination-enriched analysis of proteome

To characterise the changes observed in western blot analysis of infected samples, whole cell lysates were enriched for ubiquitin proteins using Ubiquilin pull down. A total of 3224 proteins were identified using label free mass spectroscopy (all detectable proteins), which was filtered to 1418 proteins identified against the current Uniprot proteome for Atlantic Salmon (UP000087266). Principle component analysis of 1418 proteins shows a lack of clustering by infection, with samples primarily split by time along PC1 (Figure 2A). As the focus of this research is the host response to infection, viral proteins identified in proteomics data were excluded from principal component analysis.

**Figure 2:**
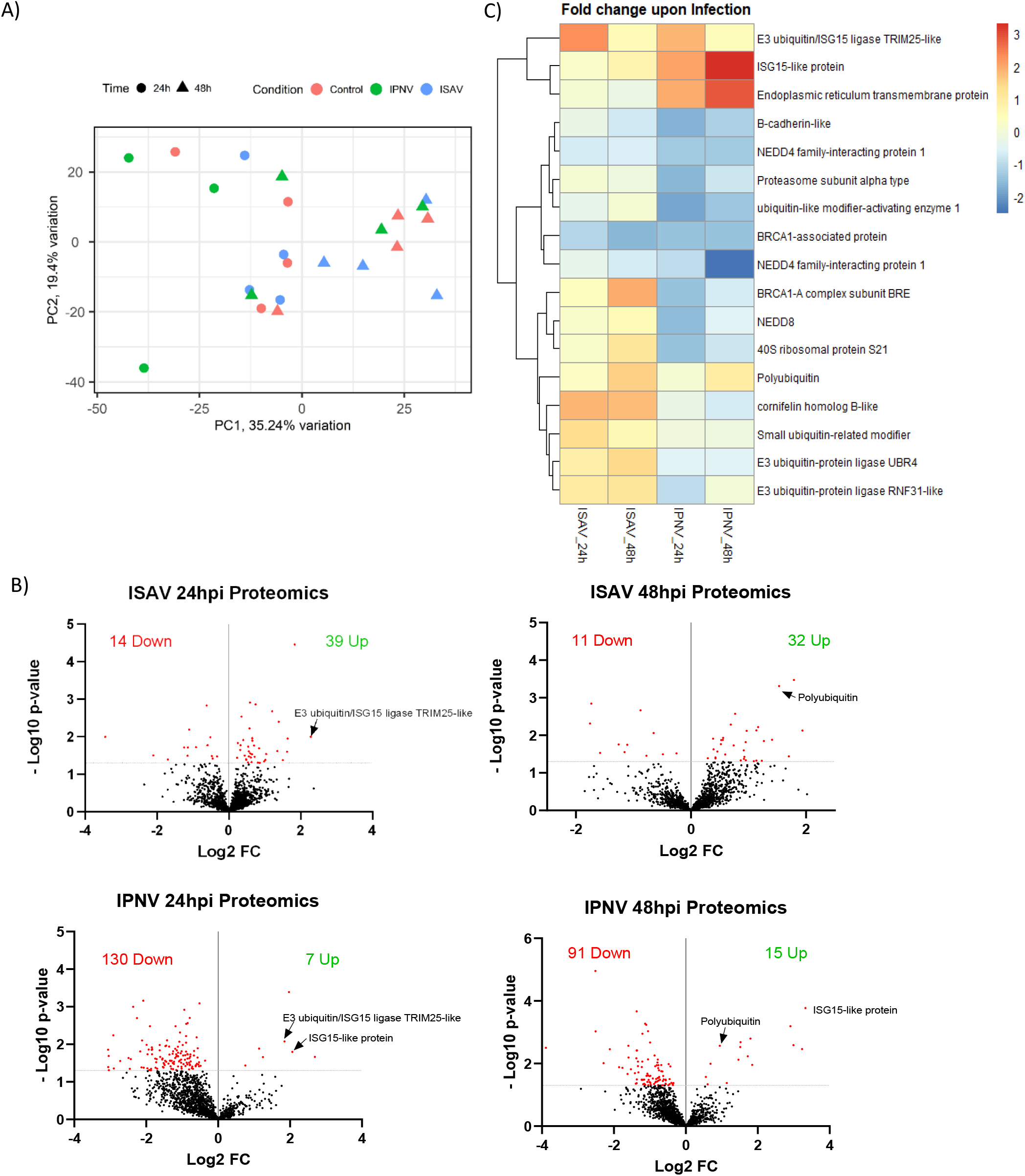
Proteomics of ISAV and IPNV infection. A) Principal component analysis of host proteins in response to infection B) Volcano plots of host proteins upon infection, significance threshold (padj<0.05) indicated by dotted line. The total number of significant proteins increased/decreased upon infection is displayed. Colours indicate Log2FC.

ISAV infection induces an increase in the abundance of ubiquitinated and ubiquitin-associated proteins, with 74% of differentially abundant proteins increasing in abundance for both 24- and 48-hour time points (Figure 2B). Conversely, a reduction in the abundance of ubiquitinated / ubiquitin-associated proteins was observed after IPNV infection, with 95% and 86% of significantly differently abundant proteins reducing at 24 and 48 hours, respectively. The number of differentially abundant proteins during ISAV infection (53 at 24 hours, 43 at 48 hours), was less than that of IPNV (137 at 24 hours, 106 at 48 hours). Although both viruses exhibit similar modes of action, the immune response involving ubiquitination in ISAV and IPNV infections is markedly different.

Multiple post translational modifying proteins are differentially abundant upon both ISAV and IPNV infection (Figure 2C). Ubiquitination pathway proteins that increase in abundance upon ISAV infection include E3 ligases UBR4 (2.7 fold change), RNF31-like protein (2.2 fold change) and Ubiquitin/ISG15 ligase TRIM25-like (4.9 fold change) – in addition to polyubiquitin (2.9 fold change), and small ubiquitin-related modifier (2.5 fold change). Post-translational proteins reducing in abundance include BRCA1-associated proteins (0.3 fold change), and Nedd4 family interacting protein 1 (0.6 fold change).

IPNV infection induced an increase in E3 TRIM25-like ligase (3.6 fold change), ISG15-like protein (10 fold change), Poly-ubiquitin (1.9 fold change) and Endoplasmic reticulum transmembrane protein (7.5 fold change). The majority of differentially abundant proteins upon IPNV are reduced, including B-Cadherin-like protein (0.3 fold change), Nedd4 family interacting protein (0.2 fold change), proteasome subunits (0.3 fold change), NEDD8 (0.4 Fold change), and multiple ribosomal proteins (0.4 fold change).

GO term enrichment analysis of differently abundant proteins for ISAV infection (Figure 3A) revealed significant enrichment of components of the translational and post translational pathway; Ribosomal pathways (>30 fold enrichment) and protein processing in the endoplasmic reticulum (>10 fold enrichment). GO enrichment analysis of IPNV infected cells (Figure 3B) demonstrates a larger response to the virus, with protein/amino acid metabolism and the proteasome being the main enriched pathways. This suggests a proteome reprogramming in response to infection which is regulated by the ubiquitin pathway. Proteasome-associated genes were enriched >60 fold, in addition to downstream metabolic degradation processes of amino acids (e.g. Beta alanine metabolism, >20 fold enrichment), and amino acid functional group metabolism (e.g. Butanoate and Propanoate metabolism, >20 fold enrichment). Whilst the most enriched terms are degradation and metabolism pathways, some biosynthesis pathways – ribosome and amino acid biosynthesis – are also enriched.

**Figure 3:**
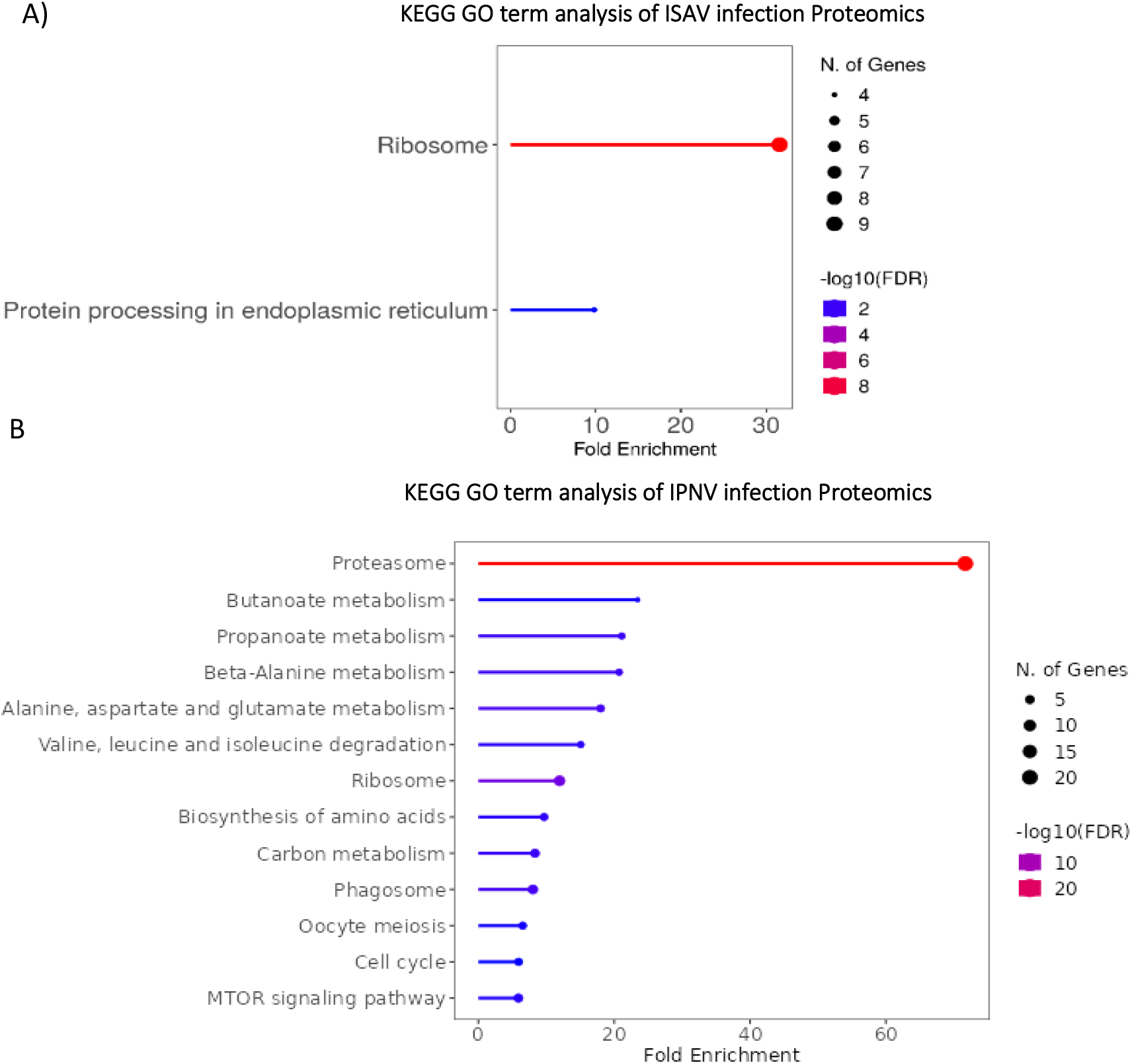
GO term analysis of infection proteomics. A) GO Term enrichment analysis of ISAV infection, 48 hours post infection, B) GO Term enrichment analysis of IPNV infection, 48 hours post infection. Go term analysis performed with R SHINYGO^29^.

### Transcriptomic reprogramming in response to ISAV and IPNV infections

RNA sequencing of infected and control samples was performed to reveal the transcriptomic response induced by the changes in the ubiquitination state. Principle component analysis of RNA sequencing data showed distinct clustering of experimental groups, with the exception of Control 24-hour and ISAV 24-hour sample groups (Figure 4B), which clustered together, suggesting a mild early response to ISAV.

**Figure 4:**
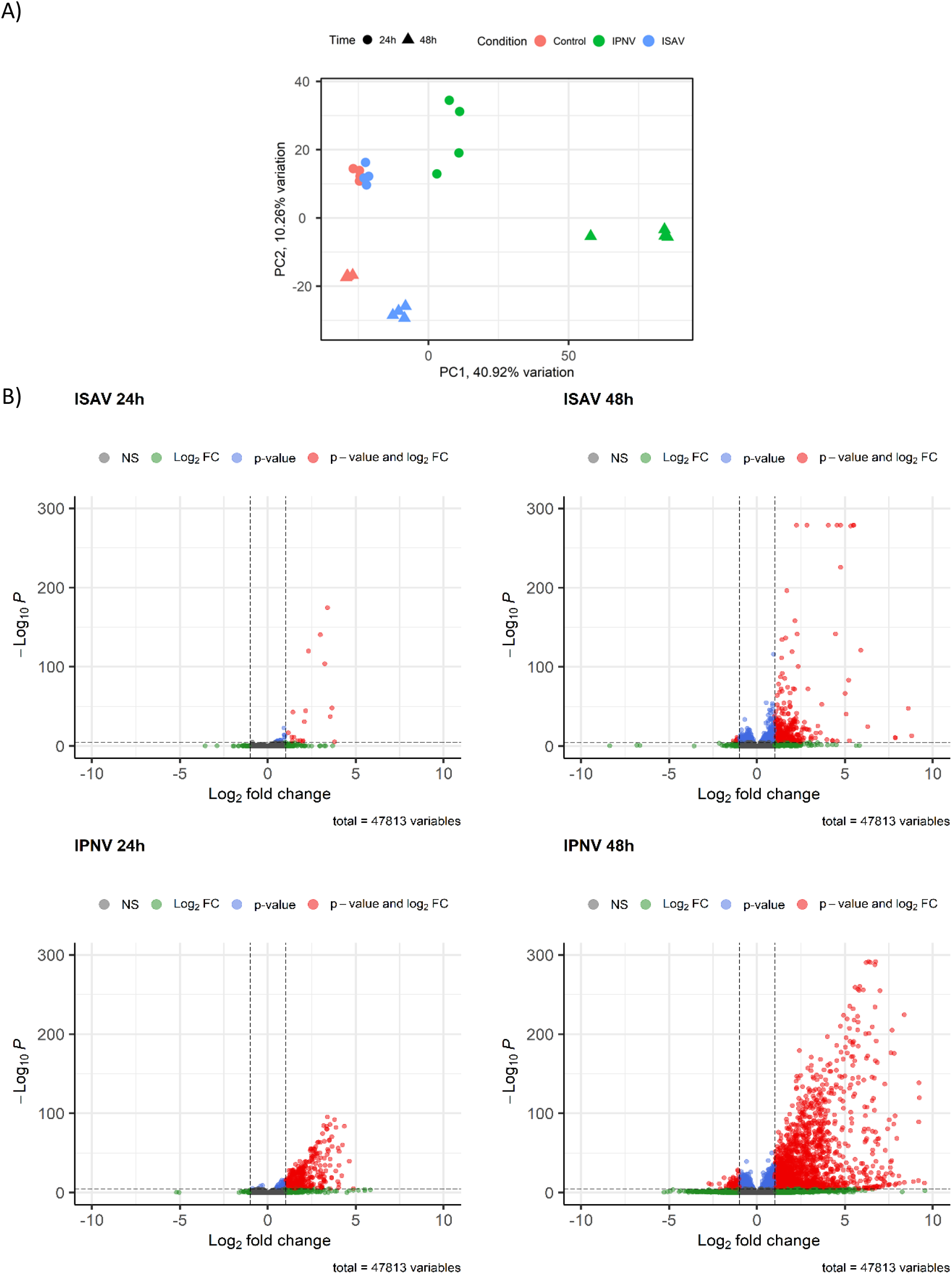
Transcriptomics of ISAV and IPNV infection. A) Principal component analysis of host transcripts in response to infection cut-off values p value <10e-6, 0.5 Log2FC B) Volcano plots of transcripts upon infection, significance threshold (padj <0.05, Log2FC >1) indicated by dotted line.

Indeed, ISAV infection resulted in a limited transcriptomic response at 24 hours, with only 36 genes significantly upregulated (p adj <0.05, FC >2; Figure 4B). Upregulated genes of note include map3k8; a regulator of the TNFα pathway, and finTRIM family member 14; a member of the expanded subfamily of TRIM proteins in fish (Figure 5A). GO term analysis of transcriptomics of ISAV infection at 24 hours produced few relevant enriched GO terms (data not shown). An increased transcriptomic response was observed 48 hours post ISAV infection, with 647 upregulated genes (padj <0.05, FC >2) (Figure 4B). Amongst the upregulated genes are classical markers of innate interferon antiviral response, best exemplified by RSAD2 (5.7 fold change), interferon alpha 1 (6.4 fold change), RIG-I (ddx58) (3.1 fold change), LGP2(dhx58) (4.8 fold change), IFIT8 (3.6 fold change), IFIT9 (4.4 fold change), IFIT10 (5.0 fold change), IRF7 (5.3 fold change), and Mx2 (4.4 fold change) (Figure 5). Many members of the conserved (TRIM) and novel (FinTRIM and Blood Thirsty) TRIM gene family are significantly upregulated in response to infection; 5 types of TRIM genes (*trim35, trim25, trim3b, trim69* and *trim16*), 8 different FinTRIM genes (*finTRIM72, finTRIM12, finTRIM66, finTRIM82, finTRIM67, finTRIM13, finTRIM14* and *finTRIM16*) and two blood thirsty gene family members (*bty18, bty4*) are upregulated. Two *Trim25* genes were upregulated in response to ISAV infection mapped to chromosomes 2 and 12, reflecting the paralogues arising from salmonid whole genome duplication^35^, yet only ssa12 *trim25* is significantly upregulated with a fold change greater than 2-fold. *Trim25* paralogue located on chromosome 12 is upregulated only 1.4-fold (padj= 0.004882), whilst *trim25* paralogue on chromosome 2 is upregulated with a greater magnitude (2.1-fold) and significance (padj= 1.80E-11).

**Figure 5:**
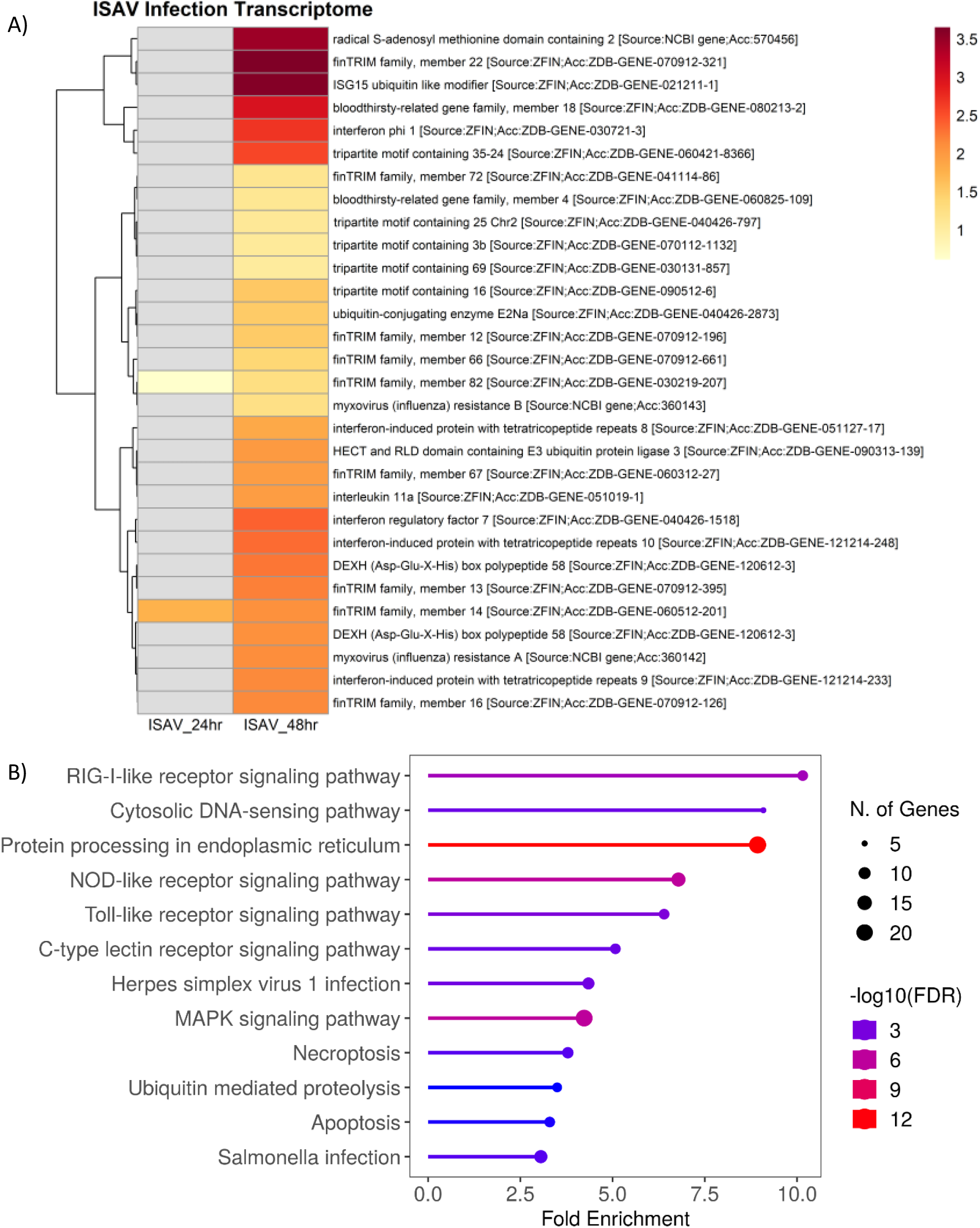
*Transcriptomics of ISAV infection in SHK-1 cells. A) Heat map of significantly differently expressed genes relating to infection and ubiquitination. Colors represent Log2 Fold change. Non-significant values are displayed in grey. B*) Enriched GO term analysis of significantly upregulated transcripts (p value <0.05, FC >2) from ISAV infection at 48 hours post infection. GO Term analysis was performed on closest homologs of salmon genes in model species; Zebrafish, using R SHINY GO^29^.

GO term analysis of upregulated genes (Figure 6A) demonstrates clear activation of the innate immune pathogen recognition receptor pathways (PRR), including RIG-I-like receptor signalling (RLR – see annotated KEGG pathway-Figure 7A), Cytosolic DNA sensing, NOD-like receptor signalling (NLR), Toll-like receptor signalling (TLR) and C – type lectin receptor signalling (CTLR). Multiple components of the protein production pathways are enriched including protein processing in the endoplasmic reticulum and ubiquitin mediated proteolysis, the latter GO term being consistent with the global upregulation of ubiquitin signalling observed in the western blot (Figure 1).

**Figure 6:**
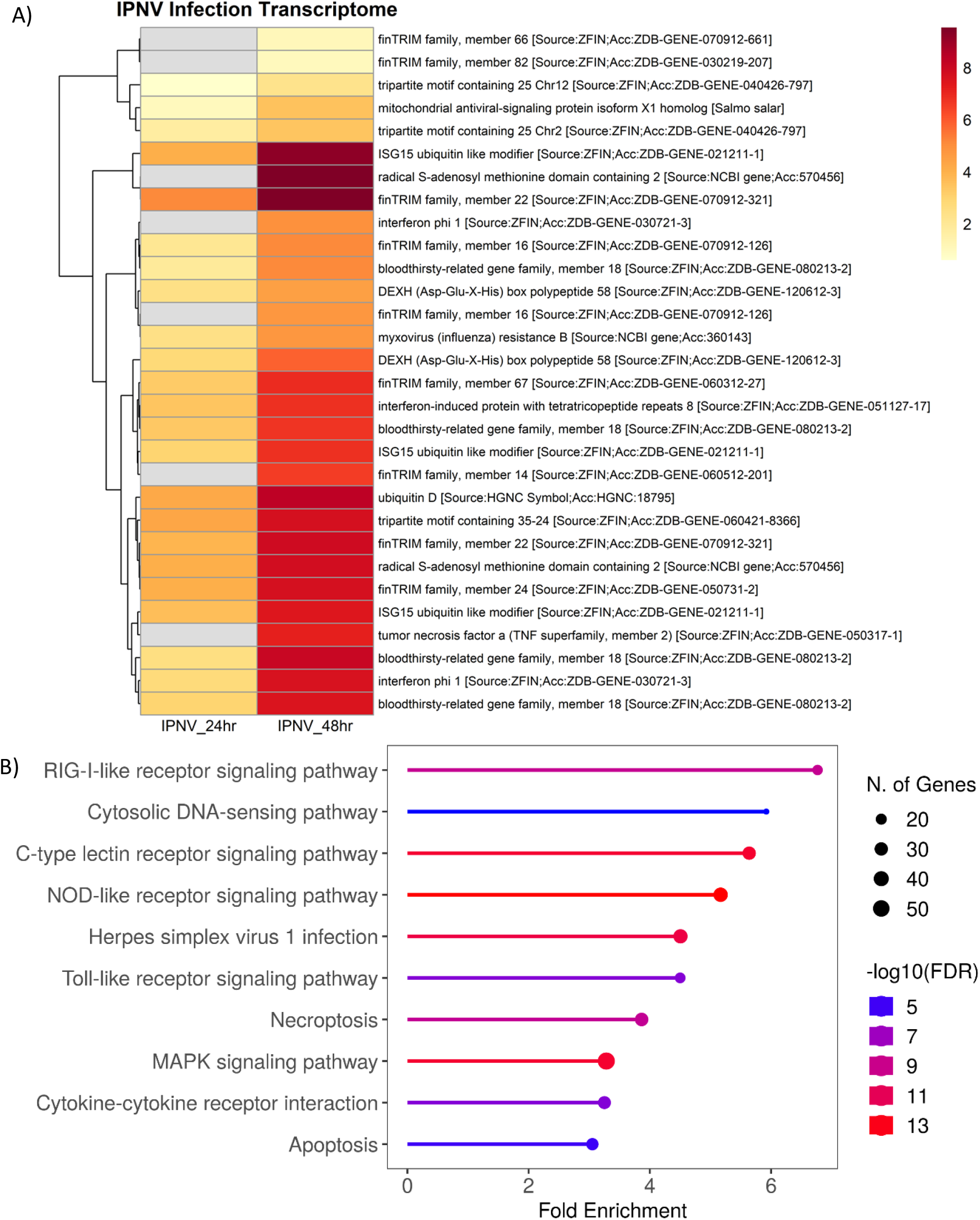
*Transcriptomics of IPNV infection in SHK-1 cells. A) Heat map of significantly differently expressed genes relating to infection and ubiquitination. Colours represent Log2 Fold change. Non-significant values are displayed in grey.B*) Enriched GO term analysis of significantly upregulated transcripts (p value <0.05, FC >2) from ISAV infection at 48 hours post infection. GO Term analysis was performed on closest homologs of salmon genes in model species; Zebrafish, using R SHINY GO^29^.

**Figure 7:**
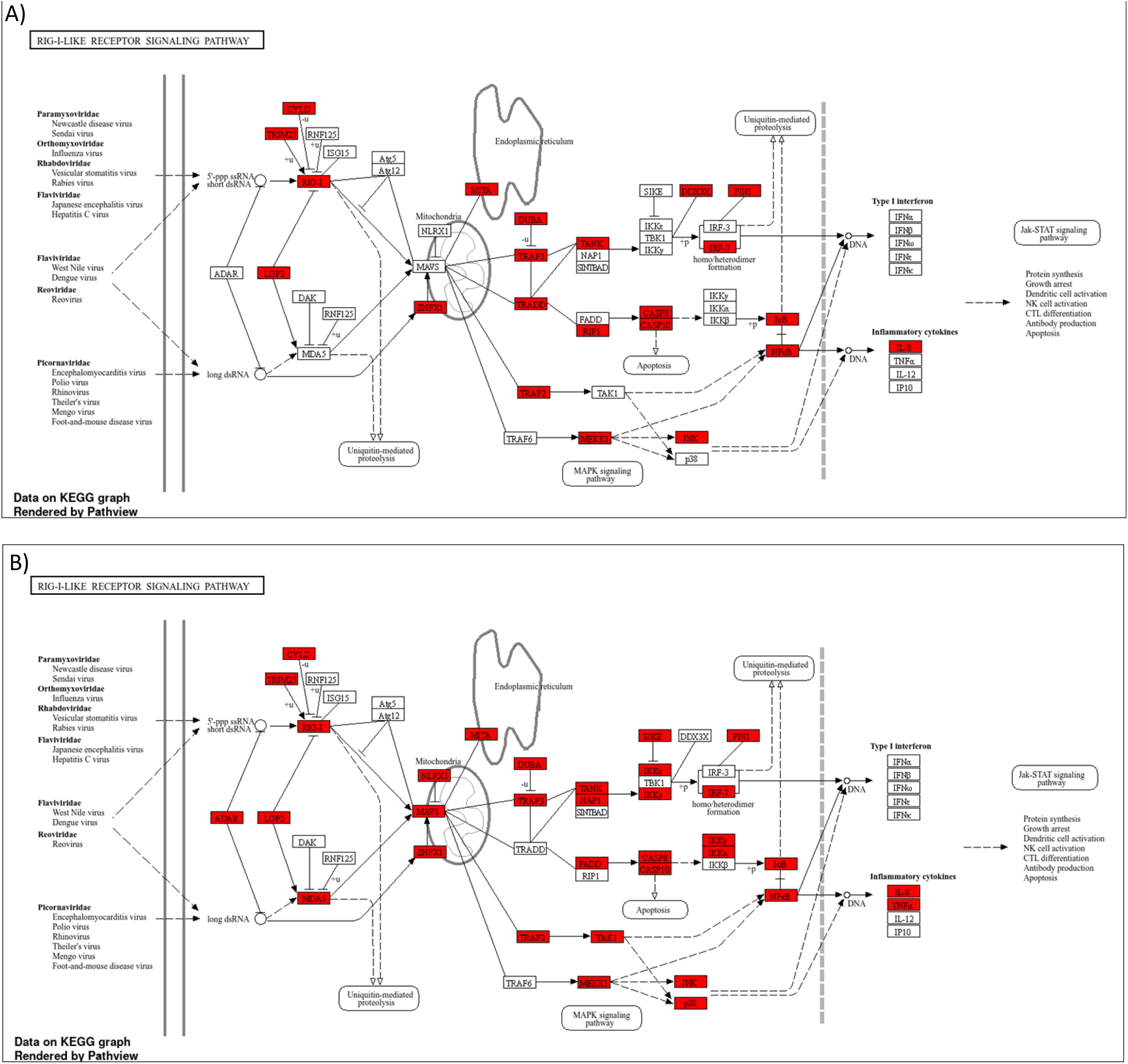
KEGG pathway annotation of zebrafish genes homologous to Atlantic Salmon genes upregulated in the RIG-LIKE receptor pathway in response to ISAV (A) and IPNV (B) Infection. Upregulated genes are highlighted in red. GO term analysis and KEGG pathway annotation was performed with R shiny GO^29^.

IPNV infection induced a rapid transcriptomic response with 435 upregulated genes at 24 hours, and 2418 at 48 hours post infection (Figure 4B). At 24 hours, classic markers of innate antiviral response are upregulated including rsad2, IFIT8, IFIT9, IFIT10, IFN, irf3 and multiple components on the Rig-like receptor signalling pathway; DEXH box peptide (LGP2), STAT1a and Mx genes (See annotated KEGG pathway-Figure 7B). Two members of the TRIM protein family; *trim25* and *trim107*, nine members of the expanded FinTRIM family (*finTRIM7*, f*inTRIM12*, f*inTRIM13*, f*inTRIM14*, f*inTRIM16*, f*inTRIM22*, f*inTRIM24*, f*inTRIM66* and f*inTRIM67*), and four members of the bloodthirsty TRIM-like genes (bty4, bty6, bty18 and bty26) are differentially upregulated. ISG15 (a ubiquitin like modifier) is highly upregulated, as is ubiquitin conjugating enzyme E2Na and HECT and RLD domain containing E3 ligases 3 and 4. Most of these genes are also upregulated at 48 hours post infection, and an additional 2418 genes were upregulated, including FinTRIM82 and tumour necrosis factor alpha (TNFα) (Figure 6A).

Mirroring the results observed in ISAV infection at 48 hours, IPNV infection at 48 hours induced a higher upregulation of chromosome 2 copy of *trim25* (10.7 fold change, padj= 1.04E-117), relative to the chromosome 12 copy (4.9 fold change, padj= 6.57E-68).

GO term analysis of IPNV infected cells (Figure 6B) suggests a strong reliance on the Pathogen associated molecular pattern (PAMP)/Pattern recognition receptor (PRR) signalling pathways for virus detection, with several classes of PPR including RLR, CTLR, NLR, TLR and cytosolic DNA sensing pathways highly significantly enriched.

### Ubiquitin-mediated transcriptional reprogramming in response to ISAV and IPNV infections

Weighted correlation network analysis (WGCNA) of ubiquitin-enriched proteomics and transcriptomics was utilized to identify clusters of genes putatively regulated by the ubiquitination of the proteins identified in our analyses. WGCNA was performed on a matrix combining the normalised expression and protein levels obtained in the RNA sequencing and proteomics datasets created in this study.

*Trim25* (both chromosome 2 and chromosome 12) genes were associated with a cluster containing 5 other TRIM-like genes, namely *trim33, trim37, trim69, trim71*, and *trim105*. RNA sensors RIG-I (dhx58) and MDA5 (ifih1), which are known viral double and single stranded RNA sensors respectively, and IFN pathway activators were associated with this cluster, suggesting a potential activation pathway mechanism. Additionally, ubiquitin conjugating enzyme E2N (*ube2n*), a known interactor with TRIM25 in human, is present in this node suggesting a putative conserved E2/E3 ligase pair. A well-known antiviral interferon stimulated gene, Protein Kinase R (PKR, *eif2ak2*) is also associated with this cluster along with neddylation genes Nedd4 binding protein 1 (*n4bp1*) and Nedd4 like E3 protein ligase (*nedd4l*). The proteins associated with this cluster of genes include E2 ubiquitin-conjugating enzyme (B5X3E0), proteasomal subunit beta (A0A1S3LZZ6) and a probable ubiquitin ligase (A0A1S3L7A4). GO term analysis of this cluster was performed with R Shiny, revealing significant enrichment of RIG-Like, Toll-like and NOD-like receptor signalling pathways, in addition to multiple innate immune, cell cycle and viral infection pathways (Figure 8). The differentially ubiquitinated E2 ubiquitin-conjugating enzyme and probable ubiquitin ligase represent potential regulators of the antiviral immune response in Atlantic salmon.

**Figure 8:**
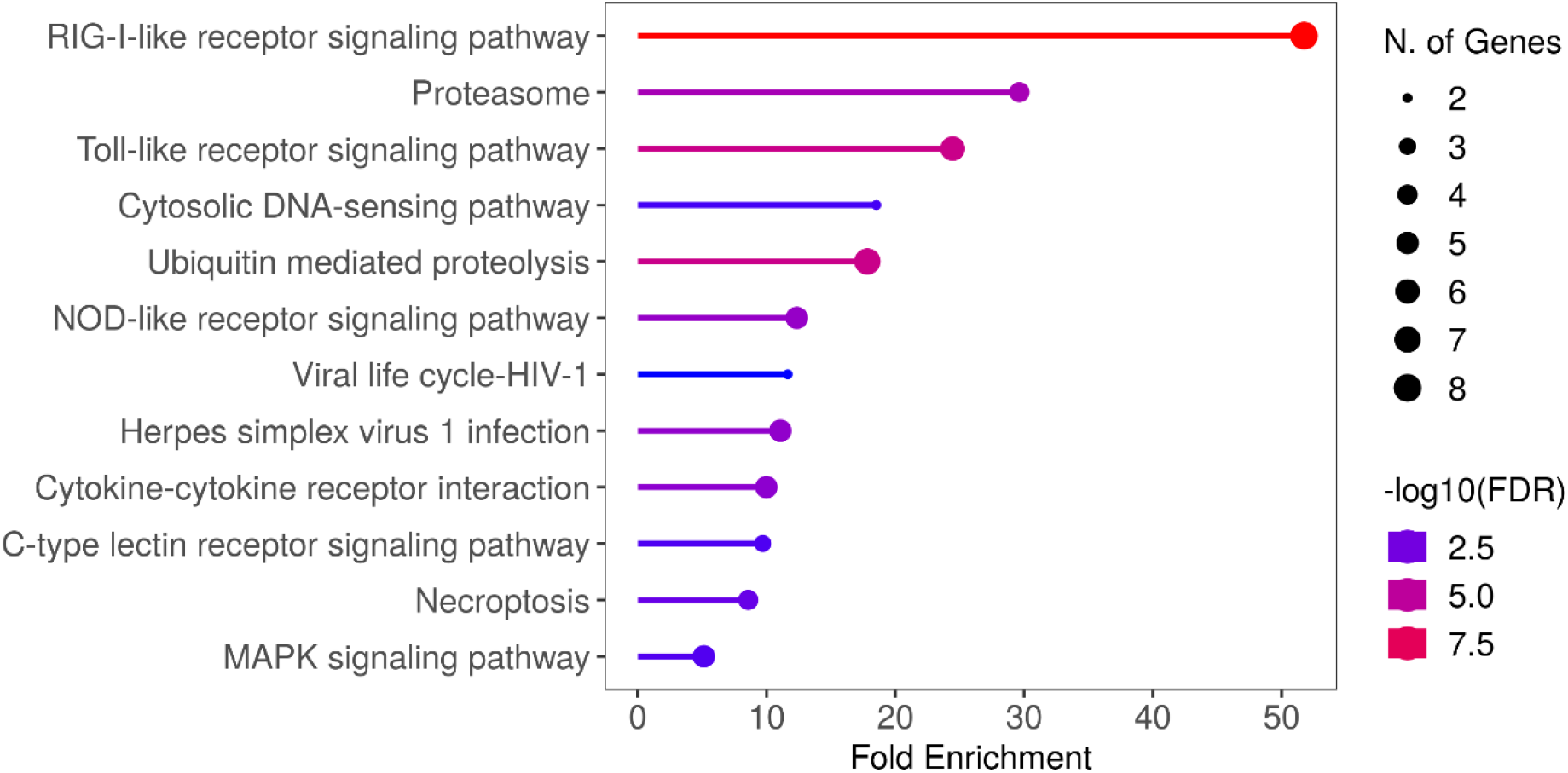
GO term analysis of trim25 containing node of WGCNA analysis. All infections and timepoints were pooled and WGCNA analysis was performed on RNA sequencing and proteomics analysis. Genes from node containing TRIM proteins (x7) was subjected to GO term analysis using Shiny Go 0.80. Zebrafish homologs of Atlantic Salmon genes were used for GO term analysis.

Poly ubiquitin is differentially abundant upon infection with both ISAV and IPNV. The network analysis reveals ubiquitin protein (B5DGL7) is associated with multiple other components of post-translational regulation, including many post-translational modification pathway proteins; TRIM containing protein 47 (A0A1S3KR00), E3 ubiquitin ligase hectd1 (A0A1S3RC95), Nedd4 family interacting protein (B9EMY3), or ISG15-like protein (Q29W12). Additionally associated with this node are multiple components of the proteasome - proteasome regulatory subunit 13 (B5DGU8), proteasome subunit beta (A0A1S3RIW7), Ubiquilin-4-like protein (A0A1S3NPQ8). Genes associated with this node also include ubiquitin ligases *trim31, trim32, trim36* and *trim55b*. This network suggests a coordinated regulation in the reprogramming of the post-translational modification / ubiquitination machinery in response to viral infection.

## DISCUSSION

In this study ubiquitin enrichment proteomics was combined with RNA sequencing to characterize the post-translational regulation of the immune response of SHK-1 cells to infection with ISAV and IPNV. ISAV and IPNV represent two RNA viruses with significant economic consequences for salmonid aquaculture, for which no prevention or treatment is completely effective. Transcriptional response to infection for both viruses has been reported for both *in vitro*^13,36^ and *in vivo*^14,37-41^ in salmonids, however investigation of post-translational modifications of proteins is very limited, despite evidence for its importance in determining the outcome of infection e.g. Neddylation pathways are associated with resistance to IPNV^42^. By combining techniques, we have gained unique insight into the virus-induced changes to the post-translational landscape of the host, whilst also measuring the impact of the changing proteome on the host transcriptome.

### Infectious salmon anaemia virus infection induces a strong post-translational response prior to a transcriptomic response

The response of SHK-1 cells to ISAV is slower than that to IPNV. While the replication time of ISAV is currently undefined, the muted transcriptional response observed at 24 hours followed by a greater response at 48 hours is consistent with the 24 hour lag phase proposed by Falk et al (1997)^43^, and Gervais et al (2023) where ISAV infected SHK-1 cells were found to clustered with control group samples at 24 hours^44^. However, our data suggests that during early infection the cell is undergoing a post-translational response, which could drive the larger transcriptomic response observed at 48 hours. ISAV infection induced a rapid increase in total ubiquitinated proteins when analysed by western blot at the earliest timepoint of 24 hours, whilst transcriptional response was limited. This suggests that ubiquitination may be important in activating/regulating the initial host response to infection. Multiple components of the protein production and processing pathways, including the ribosome, endoplasmic reticulum and ubiquitination pathway showed increased ubiquitination at 24 hours post-infection. This suggests that rather than changing in abundance these proteins are undergoing significant changes in their ubiquitination state, indicative of a post translational modification-controlled response to infection.

Differentially abundant ubiquitinated proteins of note upon ISAV infection include polyubiquitin, ISG15-like protein and TRIM25-like protein. Polyubiquitin has previously been demonstrated to be upregulated *in vivo* in response to early infection^14^, suggesting that the ubiquitination system is amongst the first to react to infection through its known activation of PRRs in fish^18^. In addition to its immune regulatory role, ubiquitin is a master regulator of protein turnover, DNA repair and cell death^45^. An increase in ubiquitin could therefore be indicative of the cytopathic effect of the virus, however cytopathic effect does not typically occur in SHK-1 cells infected with ISAV until 9 days post infection^46^.

ISG15 is a viral PAMP inducible ubiquitin homolog (40% identical, 64% conservative substitutions^47^), as previously demonstrated by its induction by the dsRNA mimic, poly I:C^48^. ISG15 has been demonstrated to regulate ubiquitination by competing for proteasome targeting ubiquitin sites in substrate proteins^49^, yet unlike ubiquitination, ISG15 conjugation does not target substrates for degradation^50^. The role of ISG15 is not well understood, but ISG15 has been suggested to perform a multitude of roles in homeostasis, including immune regulation, and is known to be transcriptionally upregulated by ISAV infection in Atlantic salmon^13^. It remains to be established whether the ISG15 protein can be ubiquitinated; hence its presence in the data presented here, or whether the Ubiquilin enrichment utilized in this study can also enrich for ISG15 due to its similarity to ubiquitin. What is clear is that ISG15 is rapidly increasing in abundance post infection, yet whether this is an anti-viral response or a pro-viral antagonism (via competing for ubiquitin-binding sites) of the immune response remains unclear.

It is also worth considering that the lag time in response to ISAV may be due to viral antagonism of immune response. It has previously been suggested that the lack of strong interferon response for ISAV is evidence of a viral evasion mechanism^51^, consistent with that seen in influenza; which belongs to the same Orthomyxoviridae family as ISAV^52^. The question that follows is whether the lack of transcriptional response observed at 24 hours is due to a heavy reliance on post-translational activation of the immune response, or are symptomatic of viral antagonism to repress host antiviral response, for which mechanisms have been described previously for ISAV^53^.

In addition to an increase in ubiquitinated proteins associated with protein production and processing pathways, GO term enrichment analysis of upregulated genes highlighted multiple pattern recognition receptor pathways upon ISAV infection. Both extracellular receptors (CLR and TLR’s) and cytosolic receptors (NLR and RLR) genes are significantly enriched, suggesting detection of viral pathogen associated molecular patterns (PAMPs) in the cytosol of the cell by NLR and RLR pathways, in addition to endosomal/extracellular sensing by TLR and CLRs.

### Conserved and fish-specific TRIM proteins play a major role in response to ISAV in Atlantic salmon

TRIM25 is a RING E3 ligase belonging to the TRIM gene family, defined by their conserved Ring domain, B-box domain and coiled coil domain (RBCC). TRIM proteins catalyse ubiquitination, ISGylation and Sumoylation of a broad range of substrates in a diverse range of cellular processes^54^. TRIM25 is a known regulator of innate antiviral immune response in model species, though its mechanism remains controversial^20,55^. TRIM25 has repeatedly been demonstrated to be induced by ISAV infection *in vitro*^13,44^ and *in vivo*^14,51^ at the transcriptomic level, however, our data represents the first evidence of regulation of TRIM25-like at the proteomic level in fish. Taken together with current literature, transcriptional and proteomic data provides clear evidence of the virus inducibility of Atlantic Salmon TRIM25 (*ssTRIM25*). However, our data suggests an unbalanced induction of the chromosome 2 copy of *ssTRIM25*. For both ISAV and IPNV infection ssa02 *ssTRIM25* is upregulated (fold change and significance) more than ssa12 *ssTRIM25* at 48 hours post infection. This expression pattern is similar to that observed in Clark et al.^56^, where only the chromosome 2 copy of trim25 (ENSSSAG00000054152) is differentially expressed in response to infection both *in vitro* and *in vivo* when stimulated with poly I:C. Taken together, this suggests some degree of either subfunctionalization or neofunctionalization, as previously described by Lien et al^35^. Further work should validate whether only chromosome 2 copy is functional or there is a different ohnolog expression pattern in different cell and tissue types, or in response to different stimuli.

Many TRIM proteins are significantly upregulated in response to ISAV infection. Boudinot et al^57^ characterised the TRIM genes in teleost fish into trim25 like genes, trim16 like genes, novel Fish trim (ftr) and ‘bloodthirsty-like’ TRIMs (btrs), of which at least one of each type is upregulated in response to ISAV infection. This confirms the importance of all four trim gene types for functional immune response in fish^54^. TRIM genes have been well characterised in model systems, where they have been demonstrated to have direct antiviral roles^26^. They can also initiate the establishment of an antiviral state by regulating cell signalling and can even form immune antagonistic roles^58^. It is clear then that the TRIM gene family that expanded rapidly at the evolution of the modern innate and adaptive immune system^58^ plays an important role in the activation and regulation of the salmon immune response to viruses. Further work should characterise and confirm the inducibility and the functions of these virus-induced E3 ubiquitin ligases.

### Infectious pancreatic necrosis virus infection induces a strong transcriptional response followed by proteasomal reprogramming

Characterisation of the post translational response to IPNV infection shows a clear deubiquitination of host proteins at both 24 and 48 hours post infection. Several protein components of the Neddylation pathway decreased in ubiquitination abundance post-infection, specifically Nedd4 family interacting protein 1 (Ndfip1), which is a regulator of HECT E3 ubiquitin protein ligases including Nedd4^59^ and a negative regulator of RIG-I-dependent immune signalling by degrading MAVS^60^. Therefore, the reduction of abundance of Ndfip1 could be associated with the upregulated transcription of the RIG-I pathway (Figure 6A). Nedd8 is also reduced in abundance upon infection with IPNV. Nedd8 has been previously associated with genetic resistance to IPNV in Atlantic salmon, where knock-out of NEDD-8 activating enzyme (*nae1*) induced a significant reduction in IPNV replication^42^. In combination with our results this highlights the important role of the neddylation post translational response to IPNV infection. The data suggests that the reduction in ubiquitination state and/or abundance of ubiquitinated Nedd8 may be indicative of increased stability of Nedd8 due to reduced proteasomal targeting via the UPS pathway.

Both ISAV and IPNV infections induce an increase in abundance of an associated ISG15-like protein, which correlates with the increased transcription of the relevant ISG15 gene. ISAV 48 hours post infection and IPNV 24 and 48 hours post infection all have a significantly upregulated ISG15 ubiquitin like modifier. ISG15 has been proposed as a negative regulator of ubiquitination by competing for substrate binding sites^61^ – this may in-part explain why ISG15 is much more abundant whilst ubiquitination state of many proteins is reduced upon IPNV infection. In addition to a possible antagonist role with ubiquitin, free ISG15 has been demonstrated to act as a negative regulator of Nedd4 in humans^62^ – this may explain the large increase of ISG15 at 48 hours post infection, paralleled with the large reduction in abundance of Nedd4 family proteins. What remains clear is that post translational modification by ubiquitin and ubiquitin-like homologs – neddylation, ISGylation and SUMOylation – are important regulators of early antiviral interferon-mediated immune response, and their regulation may temporally occur before a transcriptional response to pathogen invasion.

## CONCLUSIONS

Post translational modification of proteins by the addition of ubiquitin (and ubiquitin-like proteins) has an important role in the initiation and regulation of early antiviral immune response in Atlantic salmon in response to IPNV and, especially, ISAV. However, the responses in the ubiquitination state of the host ubiquitome to the two viruses were drastically different. For ISAV infection, ubiquitination response precedes a strong transcriptomic response to infection, whilst for IPNV there is a strong transcriptional response, however there appears to be a decrease in the majority ubiquitinated proteins post-infection. Ubiquitination of several proteins could be associated with the regulation of RIG-Like, Toll-like and NOD-like receptor signalling pathways, linking ubiquitination with the regulation of the antiviral response in this species. Ubiquitomics remains a largely understudied field in non-mammalian species, yet our results strongly suggest a key role of ubiquitin in the immune response of fish to viral infection. Further investigation of the role of this highly conserved molecule is therefore needed to provide new insights into fish immunology.

## MATERIAL AND METHODS

### Cell culture and viral Infection

SHK-1 cells (ATCC 97111106) were propagated in L15 media supplemented with 5% FBS, 40 μM of β-mercaptoethanol, 4mM of glutamine and Penicilin-Streptomicin antibiotics. At 80% confluence, cells were passaged using 0.25% of trypsin/EDTA, pelleted and split 1:3. Fresh media was added in 2:1 ratio with conditioned media. Viral stocks of ISAV and IPNV were titrated using TCID50 assays on SHK-1 to select the best concentration for viral challenge.

Viral infections (ISAV and IPNV) were performed on 80% confluent SHK-1 cells on 6 well plates. Cells were seeded then cultured overnight. Prior to inoculating cells the media was removed then replaced with L15 media with 2% FBS containing virus and incubated at 15°C for either 24 or 48 hours. A high MOI was used for each virus diluting stocks to 1/10 for ISAV and 1/100 for IPNV, alongside each inoculation a control plate without virus was also made. Cells were collected at 24 and 48 hours post-infection, using trypsin-EDTA and washed once with PBS before storing the cell pellet at -80°C either with or without TRIzol® reagent (Invitrogen™). Total RNA from samples kept in TRIzol® were extracted using Direct-zol™ RNA Microprep (Zymo research, Irvine, USA) with DNase I treatment before storing at -80°C for transcriptomic analysis. The RNA quality of each sample was checked using 4200 Tape station (Agilent) and Nanodrop, and only samples with RNA Integrity number (RIN) > 7 were used.

### Western blot

Cells were harvested at 24 and 48 hours post infection. Infected and non-infected SHK-1 cells were lysed in extraction buffer 1X PBS with 0.05% Igepal CA-630 (formerly NP-40), 0.05% Triton, 10 mM NEM, 50 μg/ml TPCK, 50 μg/ml TLCK, 0.6 mM PMSF, 100 μM MG132, 200 μM Phosphate inhibitors cocktail 1 (Sigma). Homogenates were centrifuged at 8,500g at 4°C for 15 mins to remove cellular debris. Samples were normalized for total protein content via BCA assay then subjected to electrophoresis on 8% Tris-Glycine SDS gel. Protein bands were transferred onto nitrocellulose membranes overnight at 2-8°C and blocked with 5% Non-fat dried milk (NFDM). Ubiquitinated proteins were detected by immunoblotting with anti-ubiquitin antibody (mouse anti-Ubiquitin mAb, clone P4D1, 1:4000, 1% NFDM) and visualised with Anti-Mouse IgG HRP (Cell Signalling #7076, 1:3000, 1% NFDM). Anti-pan Actin (Cell signalling #4968, 1:1000, 1% NFDM), was used as housekeeping loading control.

The remaining sample was subject to purification using HaloTag^®^ magnetic beads-bound Halo-ubiquitin (Genbank NM_053067.2, MRC PPU Reagents and Services, School of Life Sciences, University of Dundee) following the protocol described by Emmerich et al.^27^.

### Mass spectrometry of ubiquitinated proteins

Samples for proteomics were analysed by the Roslin Proteomics and Metabolomics facility. Samples were prepared for bottom-up analysis by reduction of disulphide bonds with TCEP and cysteine residues alkylated to prevent reformation with chloroacetamide. Samples were acidified with phosphoric acid and digested into peptides with trypsin via S-Trap digestion, and were analysed by LC-MS/MS using tims TOF in a data dependent acquisition. Raw mass spectral data was processed using PEAKS Studio X-Pro Software (Bioinformatics LTD). Search was performed against Uniprot Atlantic Salmon sequence database containing 47,722 entries. Data was processed either by the method described by Aguilan *et al*. ^11^ or using Perseus software^28^. Gene ontology (GO term) analysis was performed on mass spectroscopy data with R ShinyGO^29^.

### RNA sequencing

Total RNA from experimental material was used to construct polyA-enriched RNA libraries using Illumina’s TruSeq RNA Library Prep kit v2, and libraries were sequenced on an Illumna Novaseq 6000 instrument as 150 bp paired-end reads. Library construction and sequencing was performed by Novogene. The raw reads were quality filtered using Fastp v.0.21.4 ^30^. Adapter sequences were removed, reads with less than 30 bp were discarded, and low-quality bases (Phred score < 15) were filtered out. The resulting reads were pseudo-aligned against the Atlantic salmon reference transcriptome (Ensembl Ssal_v3.1) using Kallisto v0.46.1^31^. Transcript-level expression was imported into R version 4.3.2 and summarized to the gene level using tximport v1.30.0. Differential expression analysis was performed using Deseq2 v1.42.0^32^, and genes with adjusted p-values < 0.05 were considered differentially expressed. Gene Ontology (GO) enrichment analyses were performed using the web tool ShinyGO v0.77^29^ for KEGG pathway annotations. Enriched GO terms analysis was performed on the closest human homolog to the Atlantic Salmon genes that were upregulated upon infection, due to a much higher degree of annotation of the zebrafish (*Danio rerio*) genome.

### Network correlation analysis

A matrix containing normalised gene expression and protein abundance values for each sample was generated, and a weighted network correlation analysis was performed using the WGCNA package v1.72^33,34^ in R version 4.3.2. Briefly, the matrix was transformed into a weighted correlation network using a power of 3, which was then clustered into modules of highly correlated genes and proteins, allowing a minimum of 30 genes/proteins per cluster. A functional enrichment analysis was performed on specific modules using ShinyGO version 0.77^29^.

## REFERENCES

1 Food & Agriculture Organization of the United, N. The future of food and agriculture : trends and challenges. (Food and Agriculture Organization of the United Nations, 2017).

2 Béné, C. et al. Feeding 9 billion by 2050 – Putting fish back on the menu. Food Security 7, 261–274, doi:10.1007/s12571-015-0427-z (2015).

3 Janet Ranganathan, R. W., Tim Searchinger and Craig Hanson. How to Sustainably Feed 10 Billion People by 2050, in 21 Charts, 2015).

4 Houston, R. D. et al. Harnessing genomics to fast-track genetic improvement in aquaculture. Nat Rev Genet 21, 389–409, doi:10.1038/s41576-020-0227-y (2020).

5 Schubel, J. R. & Thompson, K. Farming the Sea: The Only Way to Meet Humanitys Future Food Needs. Geohealth 3, 238–244, doi:10.1029/2019GH000204 (2019).

6 Fry, J. P., Mailloux, N. A., Love, D. C., Milli, M. C. & Cao, L. Feed conversion efficiency in aquaculture: do we measure it correctly? Environmental Research Letters 13, doi:10.1088/1748-9326/aaa273 (2018).

7 MacLeod, M. J., Hasan, M. R., Robb, D. H. F. & Mamun-Ur-Rashid, M. Quantifying greenhouse gas emissions from global aquaculture. Sci Rep 10, 11679, doi:10.1038/s41598-020-68231-8 (2020).

8 Gentry, R. R. et al. Mapping the global potential for marine aquaculture. Nat Ecol Evol 1, 1317–1324, doi:10.1038/s41559-017-0257-9 (2017).

9 Gratacap, R. L., Wargelius, A., Edvardsen, R. B. & Houston, R. D. Potential of Genome Editing to Improve Aquaculture Breeding and Production. Trends Genet 35, 672–684, doi:10.1016/j.tig.2019.06.006 (2019).

10 Rimstad, E. & Mjaaland, S. Infectious salmon anaemia virus. APMIS 110, 273–282, doi:10.1034/j.1600-0463.2002.100401.x (2002).

11 Mondal, H. & Thomas, J. A review on the recent advances and application of vaccines against fish pathogens in aquaculture. Aquac Int 30, 1971–2000, doi:10.1007/s10499-022-00884-w (2022).

12 Kibenge, F. S., Godoy, M. G., Fast, M., Workenhe, S. & Kibenge, M. J. Countermeasures against viral diseases of farmed fish. Antiviral Res 95, 257–281, doi:10.1016/j.antiviral.2012.06.003 (2012).

13 Andresen, A. M. S., Boudinot, P. & Gjøen, T. Kinetics of transcriptional response against poly (I:C) and infectious salmon anemia virus (ISAV) in Atlantic salmon kidney (ASK) cell line. Developmental & Comparative Immunology 110, 103716, doi:10.1016/j.dci.2020.103716 (2020).

14 Gervais, O. et al. Transcriptomic response to ISAV infection in the gills, head kidney and spleen of resistant and susceptible Atlantic salmon. BMC Genomics 23, 775, doi:10.1186/s12864-022-09007-4 (2022).

15 Chase-Topping, M. E. et al. Impact of vaccination and selective breeding on the transmission of Infectious salmon anemia virus. Aquaculture 535, doi:ARTN 736365 10.1016/j.aquaculture.2021.736365 (2021).

16 Gomez-Casado, E., Estepa, A. & Coll, J. M. A comparative review on European-farmed finfish RNA viruses and their vaccines (vol 29, pg 2657, 2011). Vaccine 29, 3826–3826, doi:10.1016/j.vaccine.2011.03.072 (2011).

17 Liu, J., Qian, C. & Cao, X. Post-Translational Modification Control of Innate Immunity. Immunity 45, 15–30, doi:10.1016/j.immuni.2016.06.020 (2016).

18 Xiao, J., Zhong, H. J. & Feng, H. Post-translational modifications and regulations of RLR signaling molecules in cytokines-mediated response in fish. Developmental and Comparative Immunology 141, doi:ARTN 104631 10.1016/j.dci.2023.104631 (2023).

19 Zuin, A., Isasa, M. & Crosas, B. Ubiquitin signaling: extreme conservation as a source of diversity. Cells 3, 690–701, doi:10.3390/cells3030690 (2014).

20 Gack, M. U. et al. TRIM25 RING-finger E3 ubiquitin ligase is essential for RIG-I-mediated antiviral activity. Nature 446, 916–920, doi:10.1038/nature05732 (2007).

21 Inn, K. S. et al. Linear ubiquitin assembly complex negatively regulates RIG-I- and TRIM25-mediated type I interferon induction. Mol Cell 41, 354–365, doi:10.1016/j.molcel.2010.12.029 (2011).

22 Ganser-Pornillos, B. K. & Pornillos, O. Restriction of HIV-1 and other retroviruses by TRIM5. Nat Rev Microbiol 17, 546–556, doi:10.1038/s41579-019-0225-2 (2019).

23 Byk, L. A. et al. Dengue Virus Genome Uncoating Requires Ubiquitination. mBio 7, doi:10.1128/mBio.00804-16 (2016).

24 Cao, Z. et al. Ubiquitination of SARS-CoV-2 ORF7a promotes antagonism of interferon response. Cell Mol Immunol 18, 746–748, doi:10.1038/s41423-020-00603-6 (2021).

25 Zinngrebe, J., Montinaro, A., Peltzer, N. & Walczak, H. Ubiquitin in the immune system. EMBO Rep 15, 28–45, doi:10.1002/embr.201338025 (2014).

26 Rajsbaum, R., Garcia-Sastre, A. & Versteeg, G. A. TRIMmunity: The Roles of the TRIM E3-Ubiquitin Ligase Family in Innate Antiviral Immunity. J Mol Biol 426, 1265–1284, doi:10.1016/j.jmb.2013.12.005 (2014).

27 Emmerich, C. H. et al. Activation of the canonical IKK complex by K63/M1-linked hybrid ubiquitin chains. Proc Natl Acad Sci U S A 110, 15247–15252, doi:10.1073/pnas.1314715110 (2013).

28 Tyanova, S. et al. The Perseus computational platform for comprehensive analysis of (prote)omics data. Nat Methods 13, 731–740, doi:10.1038/nmeth.3901 (2016).

29 Ge, S. X., Jung, D. & Yao, R. ShinyGO: a graphical gene-set enrichment tool for animals and plants. Bioinformatics 36, 2628–2629, doi:10.1093/bioinformatics/btz931 (2019).

30 Chen, S., Zhou, Y., Chen, Y. & Gu, J. fastp: an ultra-fast all-in-one FASTQ preprocessor. Bioinformatics 34, i884–i890, doi:10.1093/bioinformatics/bty560 (2018).

31 Bray, N. L., Pimentel, H., Melsted, P. & Pachter, L. Near-optimal probabilistic RNA-seq quantification. Nat Biotechnol 34, 525–527, doi:10.1038/nbt.3519 (2016).

32 Love, M. I., Huber, W. & Anders, S. Moderated estimation of fold change and dispersion for RNA-seq data with DESeq2. Genome Biology 15, doi:ARTN 55010.1186/s13059-014-0550-8 (2014).

33 Langfelder, P. & Horvath, S. WGCNA: an R package for weighted correlation network analysis. BMC Bioinformatics 9, 559, doi:10.1186/1471-2105-9-559 (2008).

34 Langfelder, P. & Horvath, S. Fast R Functions for Robust Correlations and Hierarchical Clustering. J Stat Softw 46 (2012).

35 Lien, S. et al. The Atlantic salmon genome provides insights into rediploidization. Nature 533, 200–205, doi:10.1038/nature17164 (2016).

36 Levican-Asenjo, J., Soto-Rifo, R., Aguayo, F., Gaggero, A. & Leon, O. Salmon cells SHK-1 internalize infectious pancreatic necrosis virus by macropinocytosis. J Fish Dis 42, 1035–1046, doi:10.1111/jfd.13009 (2019).

37 Aamelfot, M., McBeath, A., Christiansen, D. H., Matejusova, I. & Falk, K. Infectious salmon anaemia virus (ISAV) mucosal infection in Atlantic salmon. Vet Res 46, 120, doi:10.1186/s13567-015-0265-1 (2015).

38 Julin, K., Johansen, L. H., Sommer, A. I. & Jorgensen, J. B. Persistent infections with infectious pancreatic necrosis virus (IPNV) of different virulence in Atlantic salmon, Salmo salar L. J Fish Dis 38, 1005–1019, doi:10.1111/jfd.12317 (2015).

39 Reyes-Lopez, F. E. et al. Differential immune gene expression profiles in susceptible and resistant full-sibling families of Atlantic salmon (Salmo salar) challenged with infectious pancreatic necrosis virus (IPNV). Dev Comp Immunol 53, 210–221, doi:10.1016/j.dci.2015.06.017 (2015).

40 Robledo, D. et al. Gene expression comparison of resistant and susceptible Atlantic salmon fry challenged with Infectious Pancreatic Necrosis virus reveals a marked contrast in immune response. BMC Genomics 17, 279, doi:10.1186/s12864-016-2600-y (2016).

41 Dopazo, C. P. The Infectious Pancreatic Necrosis Virus (IPNV) and its Virulence Determinants: What is Known and What Should be Known. Pathogens 9, doi:10.3390/pathogens9020094 (2020).

42 Pavelin, J. et al. The nedd-8 activating enzyme gene underlies genetic resistance to infectious pancreatic necrosis virus in Atlantic salmon. Genomics 113, 3842–3850, doi:10.1016/j.ygeno.2021.09.012 (2021).

43 Falk, K., Namork, E., Rimstad, E., Mjaaland, S. & Dannevig, B. H. Characterization of infectious salmon anemia virus, an orthomyxo-like virus isolated from Atlantic salmon (Salmo salar L.). J Virol 71, 9016–9023, doi:10.1128/jvi.71.12.9016-9023.1997 (1997).

44 Gervais, O. et al. Understanding host response to infectious salmon anaemia virus in an Atlantic salmon cell line using single-cell RNA sequencing. BMC Genomics 24, 161, doi:10.1186/s12864-023-09254-z (2023).

45 Komander, D. & Rape, M. The ubiquitin code. Annu Rev Biochem 81, 203–229, doi:10.1146/annurev-biochem-060310-170328 (2012).

46 Rolland, J. B., Bouchard, D., Coll, J. & Winton, J. R. Combined use of the ASK and SHK-1 cell lines to enhance the detection of infectious salmon anemia virus. J Vet Diagn Invest 17, 151–157, doi:10.1177/104063870501700209 (2005).

47 Goujon, M. et al. A new bioinformatics analysis tools framework at EMBL-EBI. Nucleic Acids Res 38, W695–699, doi:10.1093/nar/gkq313 (2010).

48 Røkenes, T. P., Larsen, R. & Robertsen, B. Atlantic salmon ISG15: Expression and conjugation to cellular proteins in response to interferon, double-stranded RNA and virus infections. Mol Immunol 44, 950–959, doi:10.1016/j.molimm.2006.03.016 (2007).

49 Kang, J. A., Kim, Y. J. & Jeon, Y. J. The diverse repertoire of ISG15: more intricate than initially thought. Exp Mol Med 54, 1779–1792, doi:10.1038/s12276-022-00872-3 (2022).

50 Perng, Y. C. & Lenschow, D. J. ISG15 in antiviral immunity and beyond. Nat Rev Microbiol 16, 423–439, doi:10.1038/s41579-018-0020-5 (2018).

51 Gervais, O. et al. Exploring genetic resistance to infectious salmon anaemia virus in Atlantic salmon by genome-wide association and RNA sequencing. BMC Genomics 22, 345, doi:10.1186/s12864-021-07671-6 (2021).

52 Gack, M. U. et al. Influenza A Virus NS1 Targets the Ubiquitin Ligase TRIM25 to Evade Recognition by the Host Viral RNA Sensor RIG-I. Cell Host Microbe 5, 439–449, doi:10.1016/j.chom.2009.04.006 (2009).

53 McBeath, A. J. et al. Identification of an interferon antagonist protein encoded by segment 7 of infectious salmon anaemia virus. Virus Res 115, 176–184, doi:10.1016/j.virusres.2005.08.005 (2006).

54 Langevin, C., Levraud, J. P. & Boudinot, P. Fish antiviral tripartite motif (TRIM) proteins. Fish Shellfish Immunol 86, 724–733, doi:10.1016/j.fsi.2018.12.008 (2019).

55 Hayman, T. J. et al. RIPLET, and not TRIM25, is required for endogenous RIG-I-dependent antiviral responses. Immunol Cell Biol 97, 840–852, doi:10.1111/imcb.12284 (2019).

56 Clark, T. C. et al. Conserved and divergent arms of the antiviral response in the duplicated genomes of salmonid fishes. Genomics 115, 110663, doi:10.1016/j.ygeno.2023.110663 (2023).

57 van der Aa, L. M. et al. A large new subset of TRIM genes highly diversified by duplication and positive selection in teleost fish. BMC Biol 7, 7, doi:10.1186/1741-7007-7-7 (2009).

58 Hage, A. & Rajsbaum, R. To TRIM or not to TRIM: the balance of host-virus interactions mediated by the ubiquitin system. J Gen Virol 100, 1641–1662, doi:10.1099/jgv.0.001341 (2019).

59 Mund, T. & Pelham, H. R. Control of the activity of WW-HECT domain E3 ubiquitin ligases by NDFIP proteins. EMBO Rep 10, 501–507, doi:10.1038/embor.2009.30 (2009).

60 Wang, Y., Tong, X. & Ye, X. Ndfip1 negatively regulates RIG-I-dependent immune signaling by enhancing E3 ligase Smurf1-mediated MAVS degradation. J Immunol 189, 5304–5313, doi:10.4049/jimmunol.1201445 (2012).

61 Fan, J. B. et al. Identification and characterization of a novel ISG15-ubiquitin mixed chain and its role in regulating protein homeostasis. Sci Rep 5, 12704, doi:10.1038/srep12704 (2015).

62 Malakhova, O. A. & Zhang, D. E. ISG15 inhibits Nedd4 ubiquitin E3 activity and enhances the innate antiviral response. J Biol Chem 283, 8783–8787, doi:10.1074/jbc.C800030200 (2008).

